# Heritability of cortical morphology reflects a sensory-fugal plasticity gradient

**DOI:** 10.1101/2020.11.03.366419

**Authors:** Uku Vainik, Casey Paquola, Xindi Wang, Yingqiu Zheng, Boris Bernhardt, Bratislav Misic, Alain Dagher

## Abstract

Human brain plastically adapts to environmental demands. Here, we propose that naturally occuring plasticity in certain brain areas should be reflected by higher environmental influence and therefore lower heritability of the structure of those brain areas. Mesulam’s (1998) seminal overview proposed a hierarchy of plasticity, where higher-order multimodal areas should be more plastic than lower-order sensory areas. Using microstructural and functional gradients as proxies for Mesulam’s hierarchy, we seek to test whether these gradients predict heritability of brain structure. We test this model simultaneously across multiple measures of cortical structure and microstructure derived from structural magnet resonance imaging. We also account for multiple other explanations of heritability differences, such as signal-to-noise ratio and spatial autocorrelation. We estimated heritability of brain areas using 984 participants from the Human Connectome Project. Multi-level modelling of heritability differences demonstrated that heritability is explained by both signal quality, as well as by the primary microstructural gradient. Namely, sensory areas had higher heritability and limbic/heteromodal areas had lower heritability. Given the increasing availability of genetically informed imaging data, heritability could be a quick method assess brain plasticity.

**Highlights (up to 85 chars):** Cortical areas vary in heritability. This is seen across structural measures.

Heritability differences could be explained by plasticity, topography, or noise.

We build a comprehensive model testing many explanations across 5 measures.

Heritability is explained by noise and 1^st^ structural gradient reflecting plasticity.

Heritability could be a method to study brain plasticity.

## Introduction

The human brain is plastic: over time, experience alters brain structure. This alteration can be detected at the level of cortical morphometry and white matter microstructure and connectivity. An extreme example is early-onset blindness which is associated with widely distributed changes in gray and white matter (Leporé et al., 2010). More every-day life examples include observations that people with different jobs, body size, or socio-economic status also vary in brain structure (Farah, 2017; Maguire et al., 2000; Vainik et al., 2018; Wu et al., 2020). Similarly, brain structure can change after cognitive training (Zatorre et al., 2012). However, theoretical proposals have outlined that brain areas could differ in their propensity for plasticity – that is, they differ in how they respond to typical environmental influences.

Such differences can be understood in the theoretical framework of synaptic plasticity outlined by Mesulam (1998). The brain is organised according to a synaptic hierarchy that follows a sensory-fugal gradient. This gradient spans from sensory-motor and unimodal areas interacting with the external world towards heteromodal and paralimbic areas that are increasingly dissociated from the here and now. It has been argued that this hierarchical segregation allows for the formation of multimodal abstract representations that underpin higher-order and self-generated cognition (Margulies et al., 2016; Murphy et al., 2018). Mesulam’s seminal model suggested furthermore that at the sensory synaptic level, plasticity is likely to be constrained, as “the accurate registration of new inputs necessitates a rapid return to a narrowly tuned baseline (Mesulam, 1998, p. 1023)”. At the same time, durable neuronal changes “would be highly useful at more downstream levels, where synaptic plasticity, induced by life experiences, could play a critical role in the adaptive modification of response patterns” (Mesulam, 1998, p. 1023). Therefore, blindness-induced plasticity is possible under certain circumstances, but the perceptual areas are likely to be less plastic than higher-order areas when observing sighted humans. On the other hand, environmental influences on brain structure are more likely to be present for higher-order regions.

It is likely, that such naturally-occurring variability in plasticity of brain structures could be reflected by variability of the heritability estimates of these structures. Heritability characterises, how much of the variation in a phenotype can be attributed to genetic factors. Phenotypic variance that is not explained by genetic factors is assumed to be explained by environmental factors and measurement error. In its simplest form, heritability can be estimated by correlating phenotype scores between monozygotic twins. Contemporary approaches use twin modelling for a more precise estimate (Visscher et al., 2008). Many studies have demonstrated differences in heritability coefficients of brain structures across multiple imaging measures, such as cortical thickness, surface area, functional connectivity, and myelination (T1w/T2w ratio) (Haak & Beckmann, 2019; Liu et al., 2019; Patel et al., 2018; Schmitt et al., 2008, 2019; Strike et al., 2019; Wright et al., 2002). Heritability estimates replicate across datasets (Guen et al., 2019; Strike et al., 2019) and across different heritability estimation methods (Guen et al., 2019).

These observed differences in heritability of regional brain measures have been proposed to reflect differences in plasticity (Haak & Beckmann, 2019; see discussion in Strike et al., 2019). The reason is that lower heritability likely marks greater environmental influence over the expression of a phenotype (Harden, 2021). In the behavioural domain, one example of environmental influence is lower socio-economic status. Lower socio-economic status can suppress the emergence of genetic potential for higher intelligence (Tucker-Drob & Bates, 2016) but supports the emergence of genetic potential for higher obesity (Silventoinen et al., 2019). In the brain, as per Mesulam (1998), the environmental suppression of heritability likely depends on the function of the brain area.

Here, we seek to empirically test, whether the plasticity potential, as outlined by Mesulam’s proposed hierarchy, could be reflected in heritability differences between cortical areas. To quantify plasticity potential, we use the functional connectivity and microstructural gradients (Margulies et al., 2016; Paquola et al., 2019). These gradients come from data driven dimensionality reduction of the whole brain functional connectomes (Margulies et al., 2016; Vos de Wael et al., 2020) and microstructural covariance patterns (Paquola et al., 2019). They both exhibit a general sensory-fugal organization (Mesulam, 1998) placing sensory/motor areas at one end and transmodal and paralimbic systems at the other. However, the microstructural gradient describes decreasing laminar differentiation, more in line with Mesulam’s notion of synaptic distance from external input, whereas the functional gradient depicts a transition from locally-connected sensory regions towards a default mode core with longer-rage connectivity (Valk et al., 2020).

Other indicators of variation in brain morphometry could include brain development and evolution. Hill et al. (2010) showed how postnatal brain development is non-uniform – the surface area of lateral temporal, parietal, and frontal cortex expands almost twice as much as other regions. Similar expansion profiles are also seen when comparing human to macaque brain, suggesting recent human evolution of those areas. It is possible that areas showing greater expansion during development and evolution could be more sensitive to postnatal experience, that is the demands of the environment (Buckner & Krienen, 2013; Hill et al., 2010). Cortical maps reflecting development and evolution have indeed both been related to the genetic correlation map of cortical surface area (Schmitt et al., 2019). We therefore use those brain maps as additional predictors of heritability.

Variation in heritability estimates may also reflect measurement error. For example, smaller brain regions display lower heritability (Patel et al., 2018) because they are noisier, which reduces the ability to detect genetic effects. At the same time, heritability is unrelated to test-retest reliability of brain parcels (Haak & Beckmann, 2019; Strike et al., 2019), suggesting that noise pertains to aggregation of vertices and not to variations in signal-noise-ratio across the cortex.

Another explanation of heritability differences between regions is spatial autocorrelation. It is likely that brain parcels that are physically close together have similar features (Alexander-Bloch et al., 2018; Burt et al., 2020), including heritability estimates. In geography, this is highlighted by Tobler’s first law – “everything is related to everything else, but near things are more related than distant things” (Tobler, 1970, p. 237). This law may account for descriptions of heritability differences along the antero-posterior (Y) axis in the sagittal plane (Liu et al., 2019; Patel et al., 2018). Therefore, we account for spatial autocorrelation (Burt et al., 2020; Miller, 2004).

In sum, we seek to test the link between heritability estimates and plasticity, as indicated by the sensory-fugal brain hierarchy. We focus on cortical brain structure including cortical thickness and surface area as most widely used measures of gray matter morphology (Winkler et al., 2018), intracortical T1w/T2w ratio, a proxy for intracortical myelin content (Glasser & Essen, 2011), as well as diffusion MRI based neurite imaging that is sensitive to the microstructural context within a given voxel (NODDI; Fukutomi et al., 2018). Our primary indicators of plasticity are functional and microstructural gradients, as they are theoretically linked to Mesulam’s hierarchy of plasticity. However, we will consider several alternative explanations of heritability differences, such as evolution and developmental patterns, noise, antero-posterior axis, and spatial autocorrelation.

## Methods

Data were provided by the Human Connectome Project, S1200 release, WU-Minn Consortium (Principal Investigators: David Van Essen and Kamil Ugurbil; 1U54MH091657) funded by the 16 NIH Institutes and Centers that support the NIH Blueprint for Neuroscience Research; and by the McDonnell Center for Systems Neuroscience at Washington University (David C. Van Essen et al., 2013).

Visual inspection of T1w images by X.W. excluded 26 participants with suspected intracranial arachnoid cyst. We further excluded single monozygotic twins, and people with missing data on control variables, leaving 984 individuals. This included 274 monozygotic twins, and 629 dizygotic twins or siblings, altogether nested in 341 families (162 with 2 members, 142 with 3 members, 33 with 4 members, 3 with 5 members, and 1 with 6 members), as well as 81 single individuals. Age range was 22-37, mean = 28.81, *SD* = 3.66. Education year range was 11-17, mean = 15, *SD* = 1.77. The sample included 528 females and 456 Males. 755 identified themselves as white, 122 as Black/African American, 6 as Asian /Native Hawaiian/Other Pacific islander, and 44 used either other label or were unknown. 93 identified themselves to have Hispanic ethnicity, whereas 891 did not. Ethics statement: current analysis is secondary data analysis of publicly available Human Connectome Project, all authors have been authorised to access the database.

### Data preparation

Heritability of cortical brain structure has previously been studied across multiple structural MRI imaging measures, such as cortical thickness, surface area, and ratio of T1w/T2w images (Liu et al., 2019; Patel et al., 2018; Strike et al., 2019). We further include neurite orientation dispersion and density imaging (NODDI) to capture brain microstructure (Fukutomi et al., 2018). Altogether, computed the heritability estimates of five structural MRI measures: 1) cortical thickness and 2) cortical surface area, characterising respectively radial and tangential neuronal expansion during development (Winkler et al., 2018); 3) T1w/T2w ratio reflecting myelination (Glasser & Essen, 2011); 4) intra-cellular volume fraction (ICVF) reflecting neuronal density, and 5) orientation dispersion (OD) reflecting angular heterogeneity of neurites (Fukutomi et al., 2018).

T1w and T2w images were co-registered using rigid body transformations, non-linearly registered to MNI152 space and cortical surfaces were extracted using FreeSurfer 5.3.0-HCP (Dale et al., 1999; Fischl, 2012), with minor modifications to incorporate both T1w and T2w (Glasser & Essen, 2011). Cortical surfaces in individual participants were aligned using MSMAll (E. C. Robinson et al., 2014, 2018) to the hemisphere-matched conte69 template (D. C. Van Essen et al., 2012). T1w images were divided by aligned T2w images to produce a single volumetric T1w/T2w image per subject (Glasser & Essen, 2011). Notably, this contrast nullifies inhomogeneities related to receiver coils and increases sensitivity to intracortical myelin. Cortical thickness, surface area and T1w/T2w intensity at the midsurface were estimated for each subject.

To measure the microstructural variations of each region, neurite imaging profiles (ICVF and OD) of cerebral cortices were adopted based on the HARDI dataset (Fukutomi et al., 2018). In brief, all neurite imaging profiles were first estimated voxel-by-voxel at individual-level by using Accelerated Microstructure Imaging via Convex Optimization (Daducci et al., 2015) (intrinsic free diffusivity = 1.7 × 10-3 mm2/s), and then, the profiles on 32k fs_LR surface were extracted using individual midthickness surfaces from the HCP dataset. By adopting the HCP workbench (https://www.humanconnectome.org/software/connectome-workbench), we execute: https://www.humanconnectome.org/software/workbench-command/-volume-to-surface-mapping notably, we used the option-ribbon_constrained for ribbon-constrained mapping algorithm by inner and outer surfaces.

As vertex-based estimates were likely noisy, we used the 7 network version of Schaefer-200 parcellation (Schaefer et al., 2018) on the 32k fs_LR surface (https://github.com/ThomasYeoLab/CBIG/tree/master/stable_projects/brain_parcellation/Schaefer2018_LocalGlobal/Parcellations/HCP/fslr32k/cifti), and for each measure, average values for each of the 200 regions were computed. We replicated our analysis on the 68 parcel Desikan–Killiany–Tourville (DKT) parcellation (Klein & Tourville, 2012), as this parcellation is commonly used and shared as default across large neuroimaging studies, such as ABCD and UK Biobank (Casey et al., 2018; Elliott et al., 2018).

### Brain maps

For both parcellations, we generated brain maps, providing a number for each parcel. Parcel location was determined by midpoint xyz coordinates. Parcel size was characterised by number of vertices forming the parcel. As another estimate of noise, we used parcel signal-to-noise ratio (Fukutomi et al., 2018), which was estimated as parcel mean divided by parcel’s standard deviation. This statistic was estimated for each of the MRI measures separately across the same Human Connectome participants as used for heritability estimation.

The principle microstructural and functional gradients were generated in a previous study of 100 Human Connectome Project subjects (Paquola et al., 2019). Structural gradients were derived from T1w/T2w maps while functional gradients were based on resting state fMRI connectivity. In brief, we generated 14 equivolumetric surfaces within the cortical ribbon and sampled T1w/T2w intensities along 64,984 linked vertices from the outer to the inner surface. Microstructure profile covariance matrices were generated by averaging depth-wise intensity profiles within parcels and calculating pairwise product–moment correlations, controlling for the average whole-cortex intensity profile. Resting-state functional connectivity matrices were generated by averaging preprocessed timeseries within parcels, correlating parcel-wise timeseries and converting them to z scores. Group-average microstructure profile covariance and resting-state functional connectivity matrices were independently subjected to row-wise thresholding (90%) and transformed into cosine similarity matrices. Finally, diffusion map embedding was applied to each cosine similarity matrix to identify the principle axes of microstructural or functional differentiation. In line with previous work, we focused on the first two eigenvectors, based on identifying the inflection point in the scree plot. The first functional and structural gradients (G1_FN_ and G1_MPC_) characterise the major gradient from sensation to cognition – from sensory-motor and unimodal areas to heteromodal and paralimbic areas. The second functional gradient (G2_FN_) ranges from somatosensory and auditory areas to visual areas, and the second structural gradient (G2_MPC_) ranges from somatosensory and ventral prefrontal areas to visual areas.

Additionally, we downloaded developmental and evolutionary surface expansion maps (Hill et al., 2010) from http://brainvis.wustl.edu/wiki/index.php/Sums:About and averaged values within cortical parcels.

### Analysis

Heritability of the parcel-wide features was estimated in R 3.6.3 (R Core Team, 2013) using SOLAR-Eclipse 8.4.2 (Kochunov et al., 2015) (http://www.nitrc.org/projects/se_linux; http://solar-eclipse-genetics.org/index.html) via the solarius package (Ziyatdinov et al., 2016) controlling for background variables: age, sex, race, ethnicity, and proxy brain volume. The categorical variables were recoded as dummies, with white or nonhispanic as reference category. The solarPolygenic() formula was parcel∼age+sex+black+asian+other+ethnicity+proxy brain volume. Proxy brain volume was the sum of Freesurfer variables: CorticalWhiteMatterVol + CSF + SubCortGrayVol) (Karama et al., 2011).

Product moment correlations between brain maps across parcels was presented using ggcorrplot (Kassambara, 2019). Predictors of heritability were compared with linear mixed models with crossed random effects and spatial autocorrelation using glmmTMB 1.0.2.1 (Brooks et al., 2017). Crossed random effects account for correlation of parcels across different MRI estimates. Because the same parcels are analysed with multiple structural imaging estimates, the current study has essentially a repeated measures design. In repeated measures, the subject and the measurement point are analysis factors, and they are crossed, because every subject is in every measurement point. In our current analysis, instead of participants there are individual brain parcels. Spatial autocorrelation accounts for similarity of parcels due to physical proximity. We used a random intercept model without modelling random slopes, as we were mainly interested in global effects across multiple measures and we tried to avoid making the model overtly complex. The formula for final model was:

Heritability ~ control variables + theoretical variables + exp(position + 0 | measure) + exp(position + 0 | parcel)

Control variables were number of vertices, signal-to-noise, and xyz coordinates. Theoretical variables were structural and functional gradients, developmental, and evolutionary maps. Position refers to coordinates of the parcel in MNI space. Exp stands for exponential modelling of spatial autocorrelation, as used in other brain imaging analyses (Burt et al., 2020). We applied False Discovery Rate (FDR; Benjamini & Yekutieli, 2001) correction to highlight the individual predictors in the final model.

We tested step-by step, whether it made sense to add the spatial autocorrelation, random effects, control variables, and predictor variables. The improved model fit was assessed in terms of AIC, BIC change. Here, lower is better, with a meaningful difference of ∼4 units (Burnham & Anderson, 2004). Once fixed effects, that is control and explanatory variables were introduced, improvement was assessed in terms of marginal R^2^, higher is better. Marginal R^2^ characterises the R^2^ of fixed effects (Nakagawa et al., 2017). Model fit statistics were extracted with the anova() function in R, and marginal R^2^ estimated with the performance::r2 function (Lüdecke et al., 2020). To get a sense, whether certain MRI measures are more important than others, we conducted leave-one-out cross-validation, leaving out one MRI measure and repeating the analysis. We also relied on several other useful packages, such as WriteXLS, broom, ggpmisc, vroom, patchwork, psych, data.table, synthpop, tidyverse, cowplot, magick (Aphalo & Slowikowski, 2020; Dowle et al., 2020; Hester & Wickham, 2020; Nowok et al., 2016; Ooms, 2020; Pedersen, 2020; Revelle, 2014; D. Robinson et al., 2020; Schwartz, 2020; Wickham & RStudio, 2017; Wilke & Wickham, 2016).

Data/code availability statement: individual-level data can be accessed from Human Connectome Project. We provide preprocessing scripts, analysis scripts, synthetic individual-level data, and summary statistics at https://osf.io/dwjr5/.

## Results

We found that brain parcels differ in heritability (Guen et al., 2019; Liu et al., 2019; Patel et al., 2018; Wright et al., 2002). Fig. 1a provides the average heritability estimates across the five studied MRI measures. At a glance, it seemed that primary areas tend to be more heritable than multimodal areas. The potential control and theoretical variables explaining heritability are shown in Fig. 1b. Visually, the heritability appeared to overlap with most other variables such as signal to noise, Y axis, developmental and evolutionary patterns, as well as with the hierarchical gradients. Numeric estimates of all variables, their per parcel means, and global means are presented in Tables S2-S4.

**Fig. 1a.**
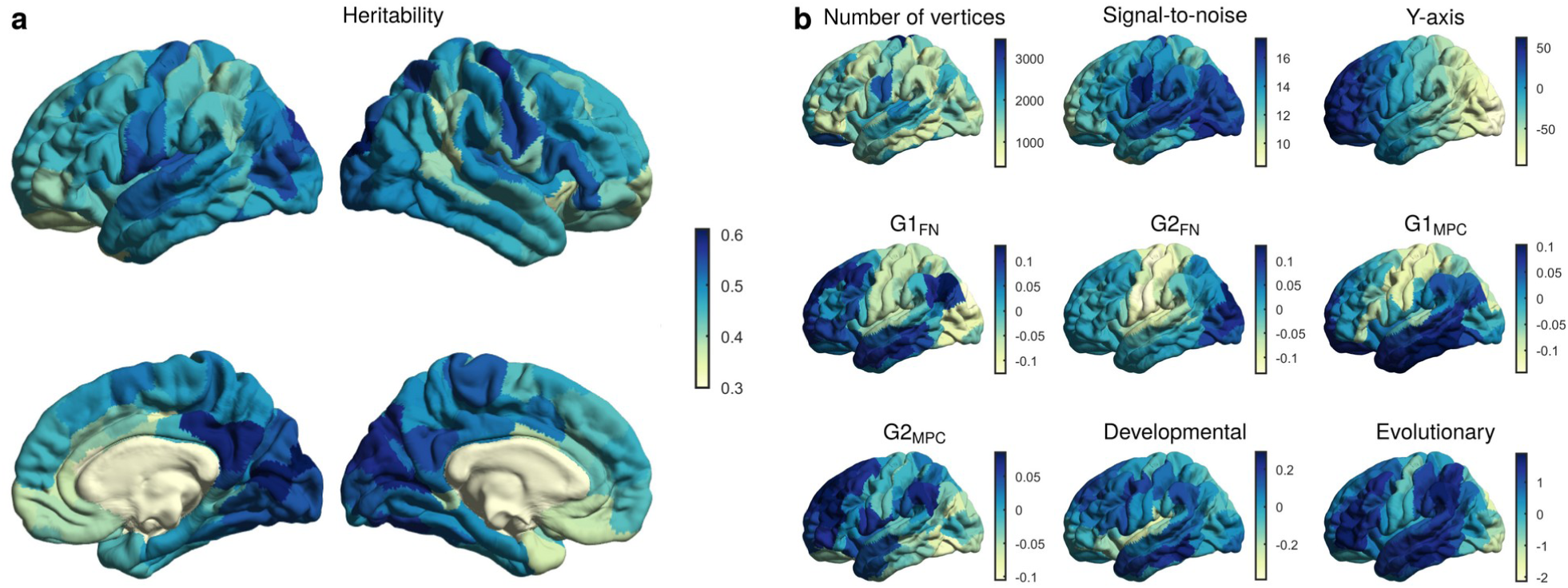
Average heritability of Schaefer-200 parcels across five measures. Top row displays lateral left and right views, and bottom row medial left and right views. Fig. 1b. Brain maps across Schaefer-200 parcels of analysed variables, using left-lateral view. We omitted X and Z axes as the visualisations are conceptually similar to Y-axis visualisation. Signal-to-noise is average signal to noise ratio across five measures. Abbreviations: G1_FN_ – first functional gradient; G2_FN_ – second functional gradient; G1_MPC_ – first structural gradient; G2_MPC_ – second structural gradient.

To quantify this impression, we estimated spatial correlations between parcels in terms of heritability of different MRI measures, control variables, and theoretical variables (Fig. 2 and 3). Control variables were number of vertices, signal-to-noise, and coordinates. Theoretical variables were structural and functional gradients, developmental, and evolutionary maps. The heritability estimates were fairly independent from each other, but they were all associated with various control variables, mostly signal-to-noise ratio, Y-axis (antero-posterior), and Z axis (left-right). In addition, heritability estimates related to structural and functional gradients, and evolution and developmental patterns. These control and theoretical variables also related to each other. This relatedness suggests that there were multiple competing explanatory mechanisms and a single model was needed to obtain the most parsimonious explanation.

**Fig 2.**
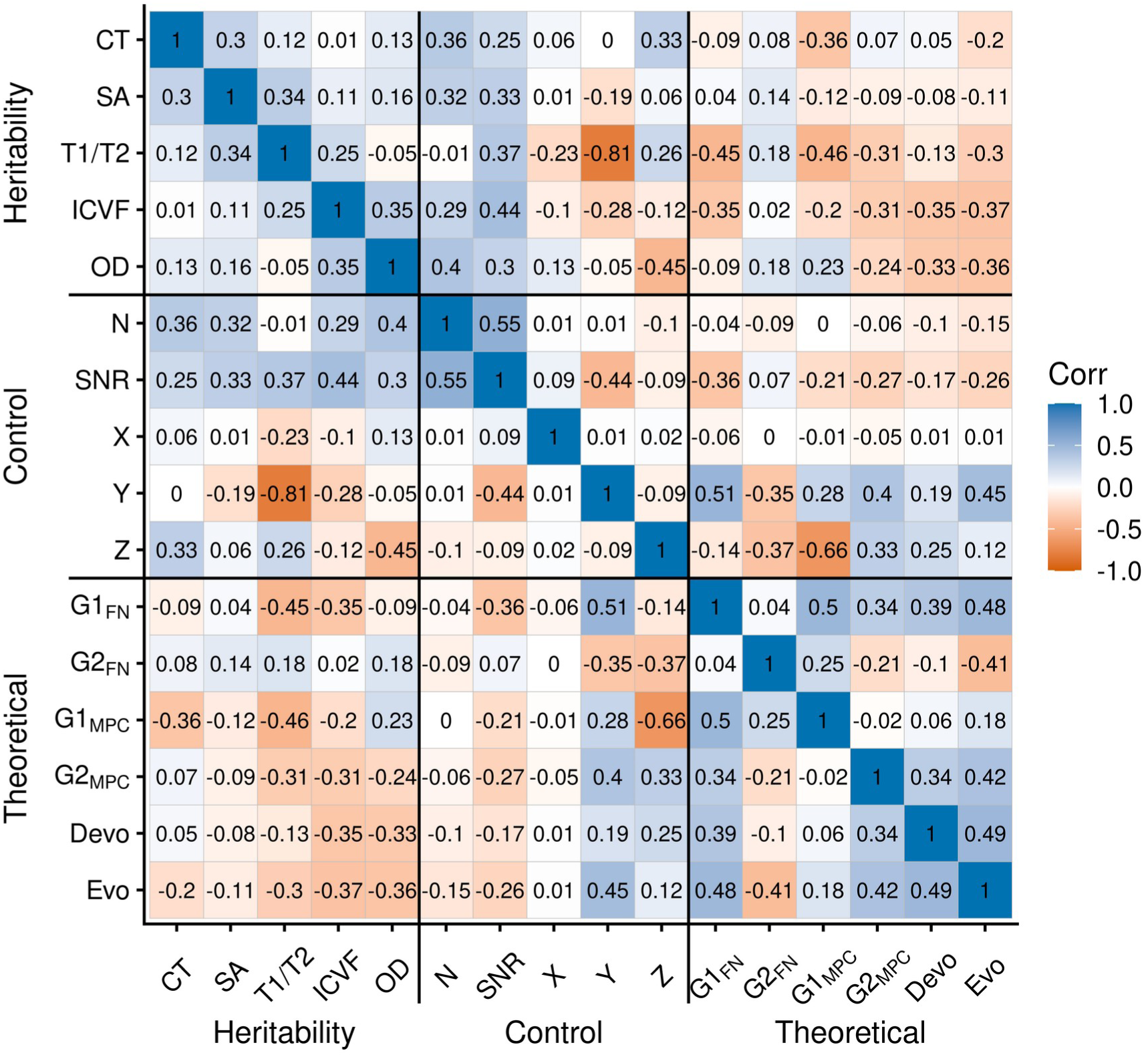
Spatial correlations between brain maps of heritability across five measures, brain maps of control variables, and brain maps of theoretical variables, using the Schaefer-200 parcellation. Abbreviations: Corr – correlation; CT – cortical thickness; Devo – developmental; Evo – evolutionary; G1_FN_ – first functional gradient; G2_FN_ – second functional gradient; G1_MPC_ – first structural gradient; G2_MPC_ – second structural gradient; ICVF – intra-cellular volume fraction; N vertices – number of vertices in a parcel; OD – orientation dispersion; SA – surface area; SNR – average signal to noise ratio across five measures; T1/T2 – T1w over T2w ratio (myelination); X,Y, Z – axis coordinates on the cortex.

**Fig. 3.**
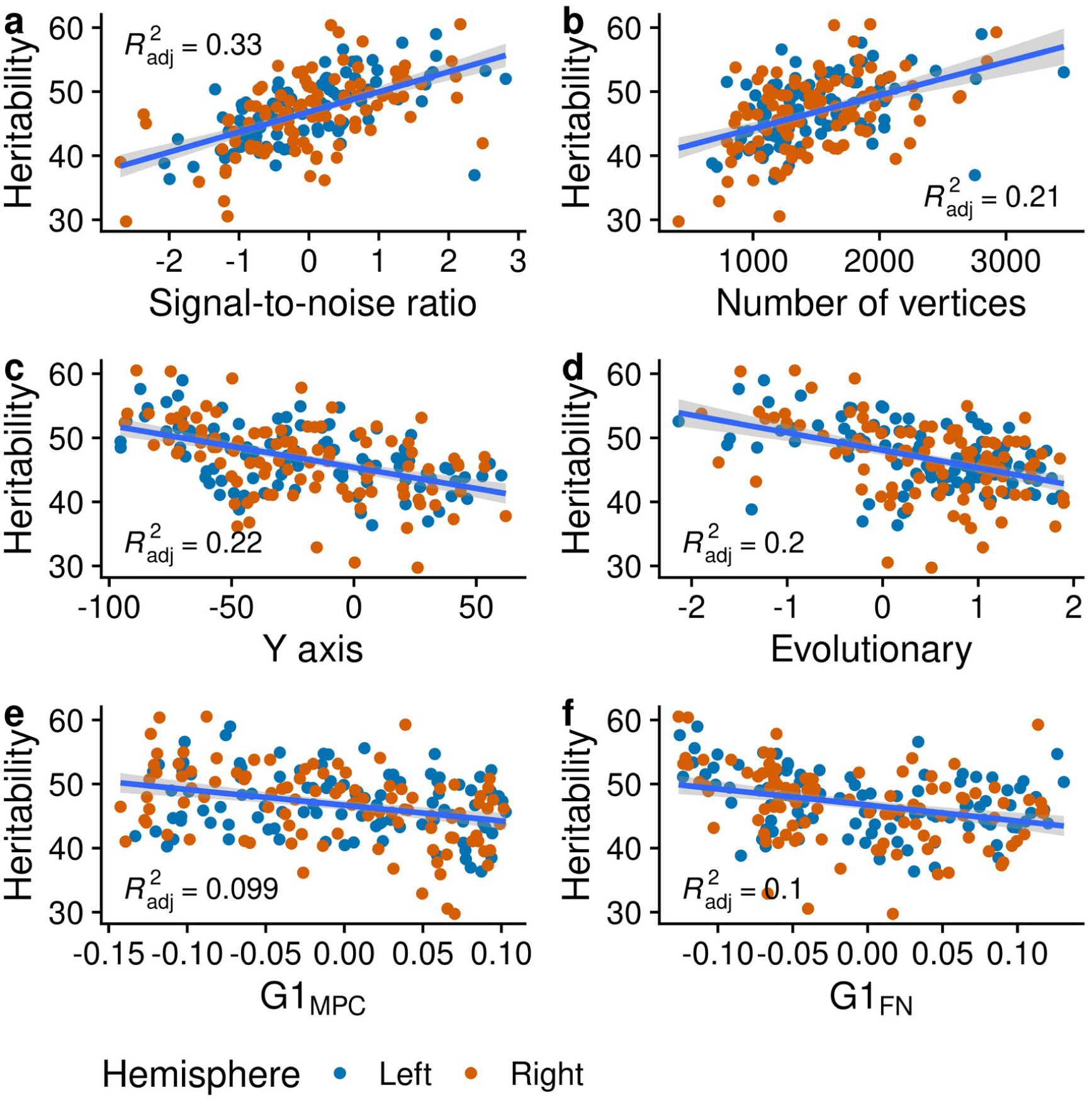
Scatterplots between mean heritability of Schaefer-200 parcels and key control and theoretical variables. Abbreviations: G1_FN_ – first functional gradient; G1_MPC_ – first structural gradient.

To find the independent factors that influence heritability estimates, we fitted a multi-level model accounting for crossed random effects for parcels across different MRI measures and parcel spatial autocorrelation. We performed a step-by-step model comparison to show that modelling spatial autocorrelation (M.auto) is better than the null model (M0), and that modelling crossed random effects further improves the model fit (M.cross.auto), in terms of lower AIC and BIC (Table 1). We tried other spatial autocorrelation modelling methods, such as Gaussian, but they were not as good (M.cross.auto.gau in Table 1). We then added control variables, which considerably improved the model in terms of BIC and AIC (M.ctl.cross.auto in Table 1). We also tried removing spatial autocorrelation. While this removal made the Y axis effect very clear (standardised estimate = 0.23, p < 0.001), the overall model fit and fixed effects’ R^2^ dropped considerably (M.ctl.mod vs M.ctl.cross.auto in Table 1), suggesting that modelling spatial autocorrelation is better than just having xyz coordinates as covariates. The final step added all discussed theoretical variables (M.ctl.theo.cross.auto in Table 1). As the model became more complex, BIC improved very little. However, the R^2^ and AIC still improved, suggesting that we are able to explain more heritability on top of control variables (delta marginal R^2^ = 0.06).

**Table 1.**
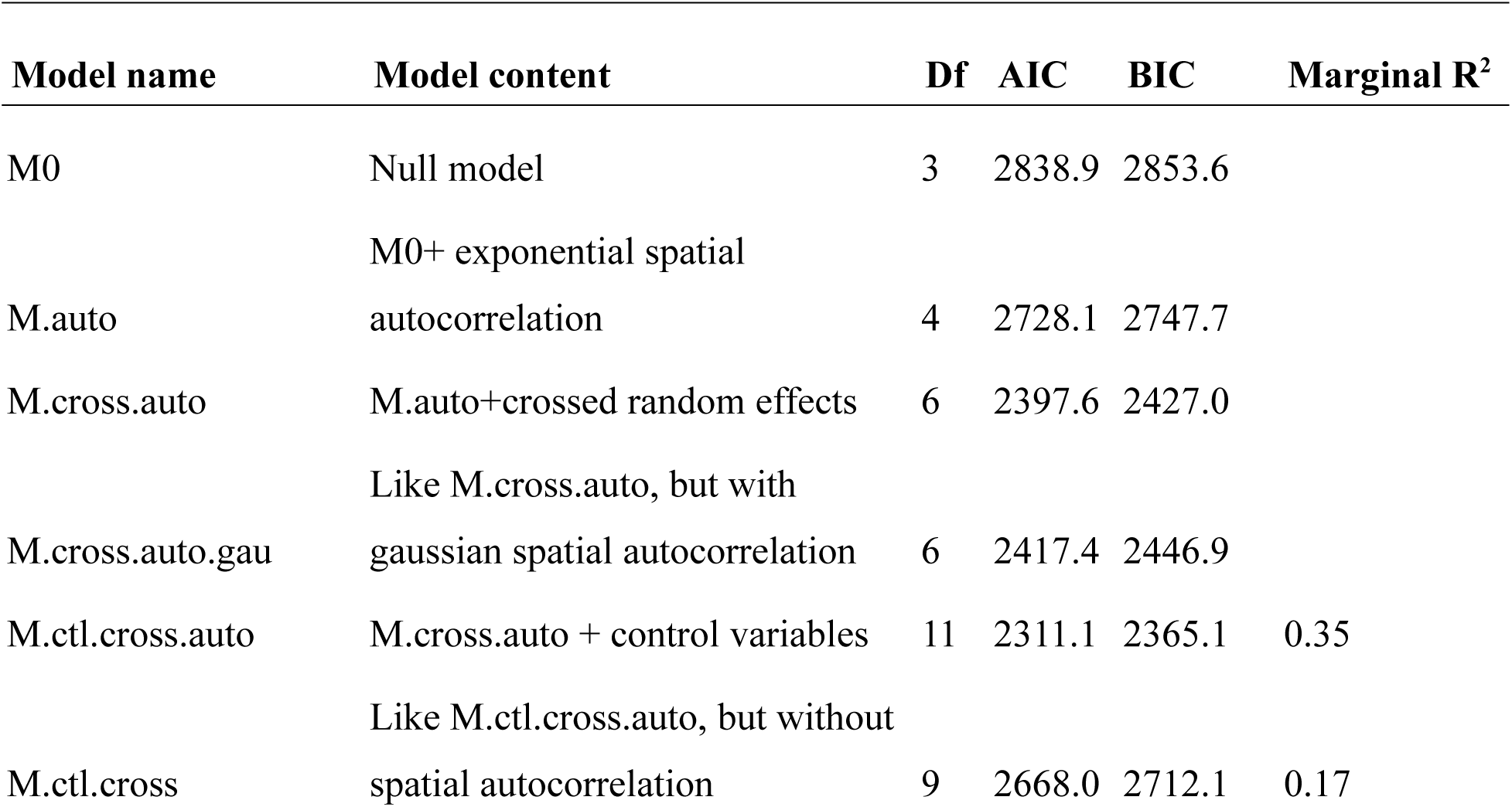
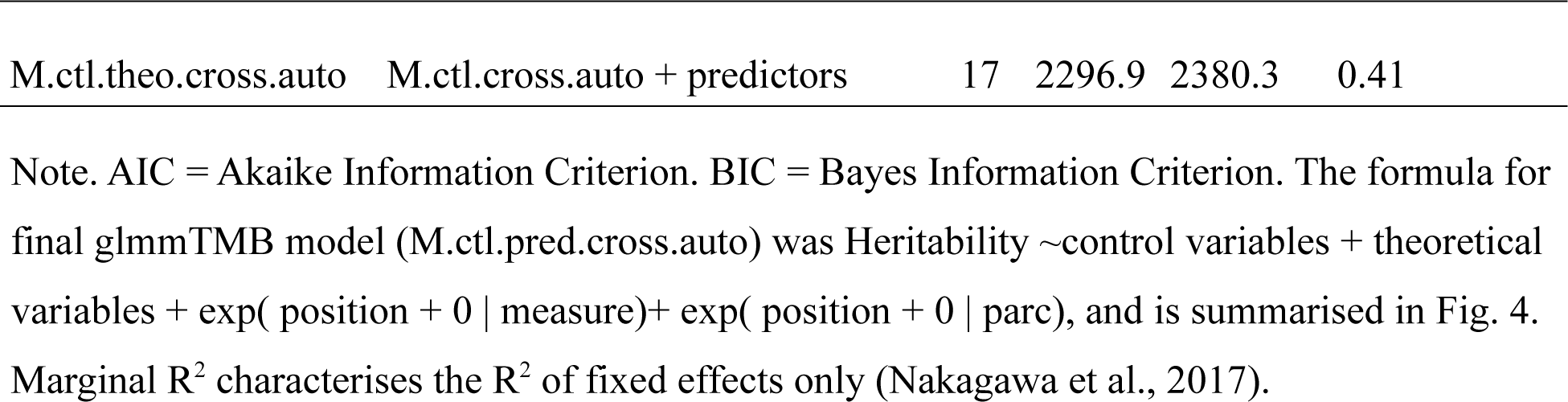
Summary of models tested.

| Model name | Model content | Df | AIC | BIC | Marginal $R^2$ |
| --- | --- | --- | --- | --- | --- |
| M0 | Null model | 3 | 2838.9 | 2853.6 |  |
| M.auto | M0+ exponential spatial autocorrelation | 4 | 2728.1 | 2747.7 |  |
| M.cross.auto | M.auto+crossed random effects | 6 | 2397.6 | 2427.0 |  |
| M.cross.auto.gau | Like M.cross.auto, but with gaussian spatial autocorrelation | 6 | 2417.4 | 2446.9 |  |
| M.ctl.cross.auto | M.cross.auto + control variables | 11 | 2311.1 | 2365.1 | 0.35 |
| M.ctl.cross | Like M.ctl.cross.auto, but without spatial autocorrelation | 9 | 2668.0 | 2712.1 | 0.17 |
| M.ctl.theo.cross.auto | M.ctl.cross.auto + predictors | 17 | 2296.9 | 2380.3 | 0.41 |

Note. AIC = Akaike Information Criterion. BIC = Bayes Information Criterion. The formula for final glmmTMB model (M.ctl.pred.cross.auto) was Heritability ∼ control variables + theoretical variables + $\exp(\text{position} + 0 \mid \text{measure}) + \exp(\text{position} + 0 \mid \text{parc})$ , and is summarised in Fig. 4. Marginal $R^2$ characterises the $R^2$ of fixed effects only (Nakagawa et al., 2017).

Fig. 4 summarises the effects of the final model. As suggested in the introduction, control variables, such as higher number of vertices per parcel and better signal to noise ratio improved heritability estimates. The MNI coordinates had low effects, as they were likely accounted for by spatial autocorrelation. Altogether, the control variables without spatial autocorrelation explained 17% and with spatial autocorrelation 35% of the variance, suggesting that control variables explain substantial part of heritability differences between parcels.

**Fig 4.**
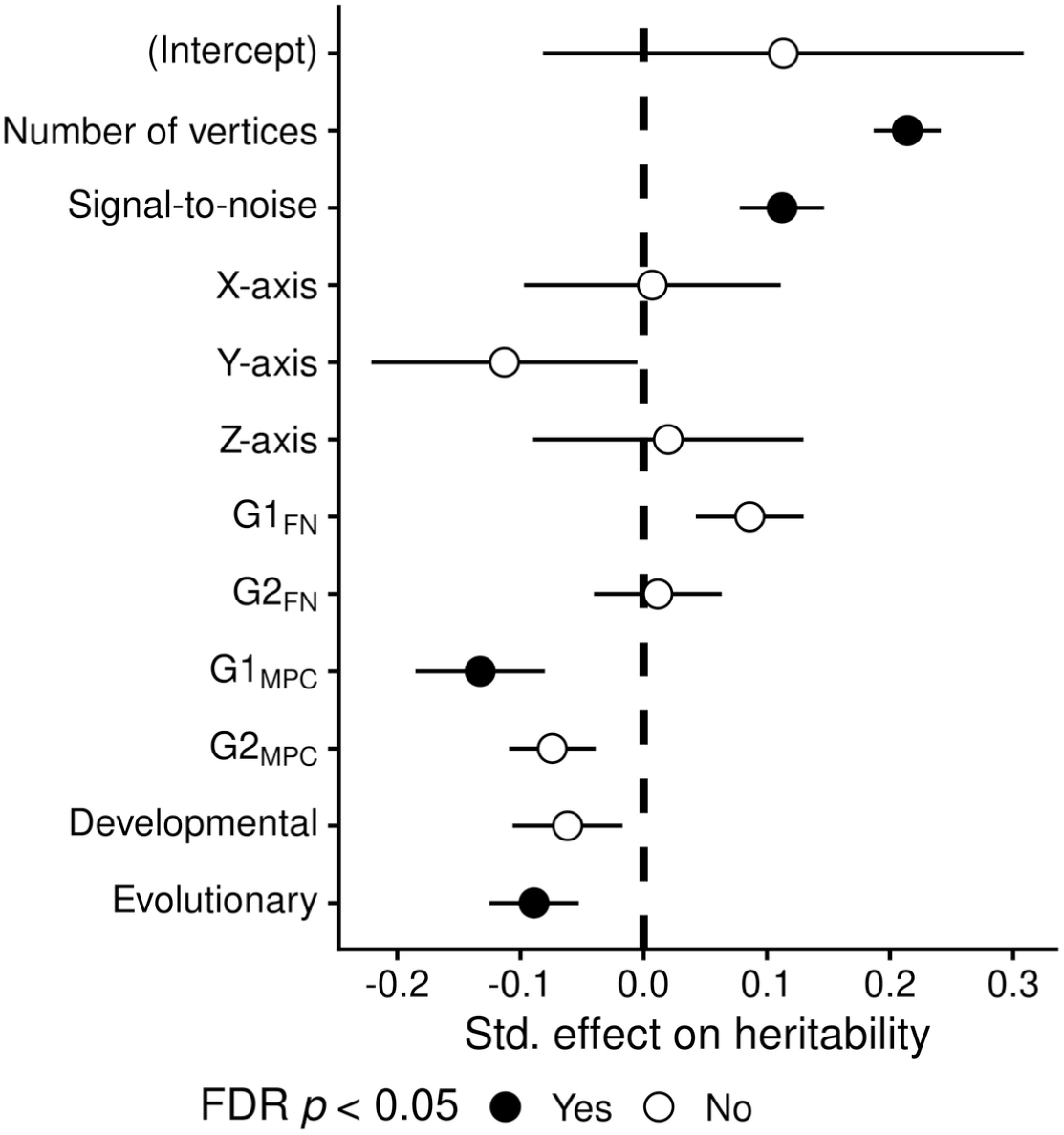
Independent effects of variables explaining heritability, based on Schaefer-200. XYZ axes are presented, but their effect is zero, as we account for spatial autocorrelation structure. See Supplementary Fig. 4S for DKT replication. Abbreviations: G1_FN_ – first functional gradient; G2_FN_ – second functional gradient; G1_MPC_ – first structural gradient; G2_MPC_ – second structural gradient. Numeric values are presented in Table S5.

The hypothesised theoretical variables increased model R^2^ to 41%. The strongest effect was a negative relation between heritability and the first microstructural gradient ranging from sensation to cognition. This supports Mesulam’s theory that higher-order structures have lower heritability due to greater plasticity (Mesulam, 1998). There was a slight additional effect of the evolutionary brain map, supporting the notion that recently evolved areas may have lower heritabilities. Heritability was also linked with G1_FN_ and G2_MPC_ at uncorrected *p*-value thresholds, but the associations became *p* > 0.05 once FDR correction was applied.

To get a sense of robustness of the results with respect to preprocessing choices, we repeated the main analysis pipeline using the DKT parcellation (supplementary Fig. S1-S4, Table S1). The DKT parcellation follows sulco-gyral folding patterns, has fewer parcels (68) and parcel sizes are more unequal. Expectedly, there is variability in heritability and in other analysed variables (Fig. S1, Tables S6-S8). As can be seen in Fig. S2-S3, the heritability of parcels is more similar to each other across MRI measures, and heritability estimates have associations with control variables and several theoretical variables.

Model fit procedure was very similar (Table S1). The final model similarly outlined the relatively strong effects of number of vertices and signal-to-noise on heritability (Fig. S4). This is expected, as DKT parcels sizes are more unequal. The first microstructural gradient had even stronger negative effect size estimate than in the Schaefer-200 parcellation (standardised estimate: -0.28 in DKT vs -0.13 in Schaefer-200). Other variables had no detectable effects on heritability.

Leave-one-measure out analysis showed that associations with larger effect size are less vulnerable to leaving out one MRI measure. The effects of control variables generally replicated across all leave-one-out iterations for both Schaefer-200 and DKT parcellations (Fig. S5 and S6, Tables S5 and S9). The effect of the first structural gradient needed the inclusion of cortical thickness, T1w/T2w and intra-cellular volume fraction to survive when using Schaefer-200 (Fig. S5), but was more robust when using DKT, as the effect was stronger (Fig. S6). The effect of evolutionary brain map only survived when orientation dispersion was excluded (Fig. S5). While DKT estimates seemed more robust, DKT modelling did not converge when cortical thickness or surface area were excluded. These results suggest that diverse MRI measures and parcellation schemes with more parcels may be needed to show heritability effects.

## Discussion

We demonstrated that heritability could reflect brain plasticity, as indexed by the microstructural sensory-fugal gradient. This association held across multiple imaging measures of gray matter morphology and microstructure and two parcellation schemes, while accounting for several other factors, such as signal-to-noise ratio of parcels and spatial proximity. Therefore, heritability could be considered as a marker for naturally occurring propensity for experience-dependent change in cortical morphometry.

The heritability patterns ultimately characterise the individual’s response to the environment. As outlined in the introduction, all brain areas can display plasticity. However, our environment usually does not require their constant reconfiguration. In response to typical demands of the environment, some areas may be more plastic than others, particularly the ones processing higher-order information (Mesulam, 1998). In particular, Mesulam suggests that it is desirable to limit plasticity in unimodal sensory areas, where faithfully transcribing information from the sense organs is paramount, while allowing experience-dependent changes in brain areas involved in cognition, emotion, planning and adaptive behaviour. This is also supported by recent findings that structure-function relationships in the brain are region-specific, with greater correspondence in primary sensory areas compared to association areas (Baum et al., 2020; Paquola et al., 2019; Preti & Van De Ville, 2019; Vázquez-Rodríguez et al., 2019). Put another way, it appears to be adaptive to favour predictable, less environmentally influenced transformations of input data in primary areas, but to allow the association cortices to be more flexible to the demands of the environment.

We have shown that heritability could be a useful metric to measure the potential plasticity of the brain. While the structural heritability patterns of the brain of contemporary free-living humans tend to be similar across datasets, this does not mean it would always have to be this way. Like plasticity, heritability is a function of the environment of the participants in the study (Visscher et al., 2008). For example, if some siblings were accidentally blind and others not, we could likely observe much higher plasticity and lower heritability in the sensory areas (Leporé et al., 2010). Luckily, this is not the case for most people.

Position on the sensory-fugal microstructural gradient was the best theoretical explainer of heritability differences. This microstructural gradient depicts a transition in myeloarchitecture across the cortex, from overall high intracortical myelin in sensory areas towards infragranular heavy microstructure profiles in paralimbic regions (Paquola et al., 2019). Intracortical myelin is thought to enhance stability by insulating fibres from making new synaptic connections (Braitenberg, 1962; Braitenberg & Schüz, 2014; Micheva et al., 2016), providing a plausible biological mechanism that links the sensory-fugal gradient to degree of plasticity. Furthermore, within the prefrontal cortex, intracortical myelin is inversely correlated with markers of plasticity, and this balance of stability/plasticity also aligns with changes in laminar differentiation (García-Cabezas et al., 2017). In line with these findings, our results further support the relationship between the cortical architecture, synaptic distance from external input and plasticity as proposed by Mesulam.

Still, the microstructural gradient is not a direct measure of plasticity. In the future, we would like to relate the heritability map to other potential indicators of plasticity, such as aerobic glycosis (Goyal et al., 2014). Among other tested brain maps, the evolutionary brain map had a detectable effect when using the Schaefer-200 parcellation. Given it did not replicate across DKT, more research is needed to understand its robustness. We suggest considering other tested brain maps in future analyses alongside the first microstructural gradient. It may also be that all those brain maps offer different nuances of the same general phenomenon, and once the plasticity-related brain maps are found, some integration of them is necessary.

Besides reflecting plasticity, heritability also depends on the noisiness of the estimates. Here, we showed heritability is higher for larger parcels with higher signal-to-noise ratio. This suggests that heritability estimates are most easily compared when parcel sizes are uniform. However, parcellations also have anatomical relevance, which may require non-uniform parcels. Nonetheless, even if parcels are more uniform, as they were in Schaefer-200 compared to DKT parcellation, variability in signal to noise ratio should still be taken into account.

Brain topography is another factor explaining variation in heritability. As parcels are not islands but relate to each other, neighbouring parcels also have similar heritability coefficients. For instance, the antero-posterior axis of brain heritability (Liu et al., 2019; Patel et al., 2018) is largely explained by spatial autocorrelation, already used for decades among geographers (Miller, 2004). While the spin test has also been proposed (Alexander-Bloch et al., 2018), that approach does not readily support multiple simultaneous predictors. Recently, spatial autocorrelation modelling has been specifically adapted for the brain – using the exponential function to model spatial autocorrelation (Burt et al., 2020), as done here.

Current analysis is limited to one sample – the Human Connectome Project composed of mostly healthy young adults. As previous comparisons of heritability across datasets and methods suggest that the heritability measures converge (Guen et al., 2019; Strike et al., 2019), we believe our results are robust. However, further analysis would allow an exploration of the generalizability of our findings to older cohorts or those affected by disease. Ideally, the noisiness of the parcel should be estimated from test-retest reliability data, as it would account for sources of measurement error beyond parcel size. Current signal-to-noise ratio may also partly capture brain plasticity, as plasticity could genuinely introduce more variance in the brain. Assessing methodological influences on our results, our main results held over two different parcellation schemes. However, more parcellations could be analysed to verify replication.

Further, it would be interesting to conduct a search for parcel size / noise trade-off (e.g., Urchs et al., 2019) to determine the optimal parcel size for heritability analysis. Possibly, a richer set of brain maps could emerge as predictors if more fine-grained parcellation is used. Similarly, it would be intriguing to widen the analysis to functional data (e.g., ArnatkeviČiūtė et al., 2020) to see if the outlined structural principles hold.

Taken together, we have shown that the primary microstructural gradient explains part of the inter-regional heritability differences of brain morphology and microstructure. This association held across multiple structural imaging measures and two different parcellation schemes, while accounting for noisiness and spatial autocorrelation of the parcels. We also outlined the general principles of using multiple brain maps to explain a patterns of interest. As genetically informed brain imaging samples become larger and more available, heritability could become an important window into brain plasticity.

