## Supplementary material for "Heritability of cortical morphology reflects a sensory-fugal plasticity gradient": Fig. S1-S6; Table S1

Data repository: https://osf.io/dwjr5/

Tables S2-S9 are available in a separate .xlsx file

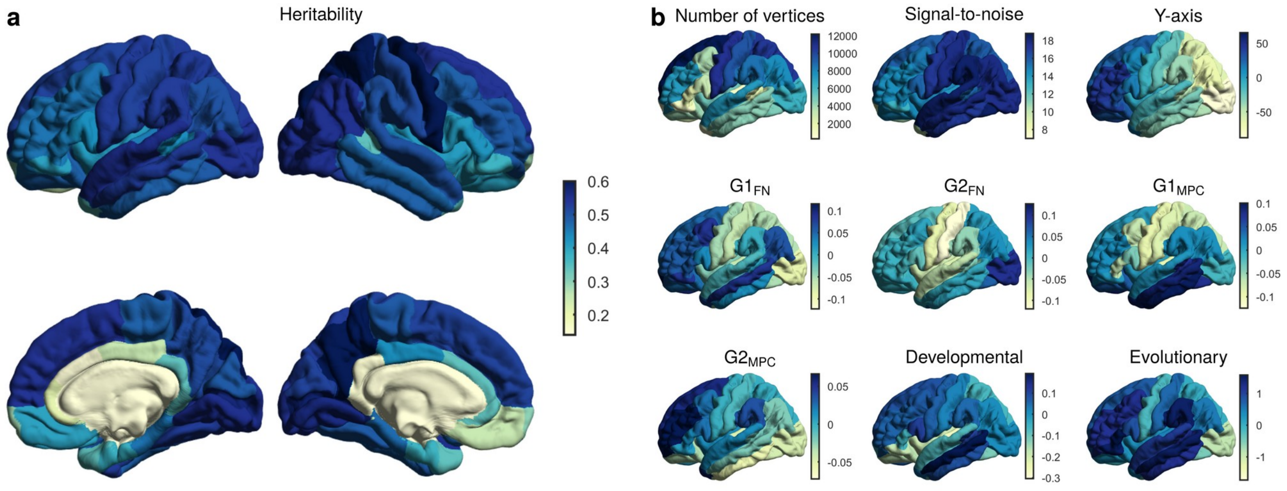


Fig. S1a. Average heritability of DKT parcels across five modalities. Top row displays lateral left and right views, and bottom row medial left and right views. Fig. S1b. Brain maps across DKT 200 parcels of analysed variables, using left-lateral view. We omitted X and Z axes as the visualisations are conceptually similar to Y-axis visualisation. Signal-to-noise is average signal to noise ratio across five modalities. Abbreviations: $G1_{FN}$ – first functional gradient; $G2_{FN}$ – second functional gradient; $G1_{MPC}$ – first structural gradient; $G2_{MPC}$ – second structural gradient.

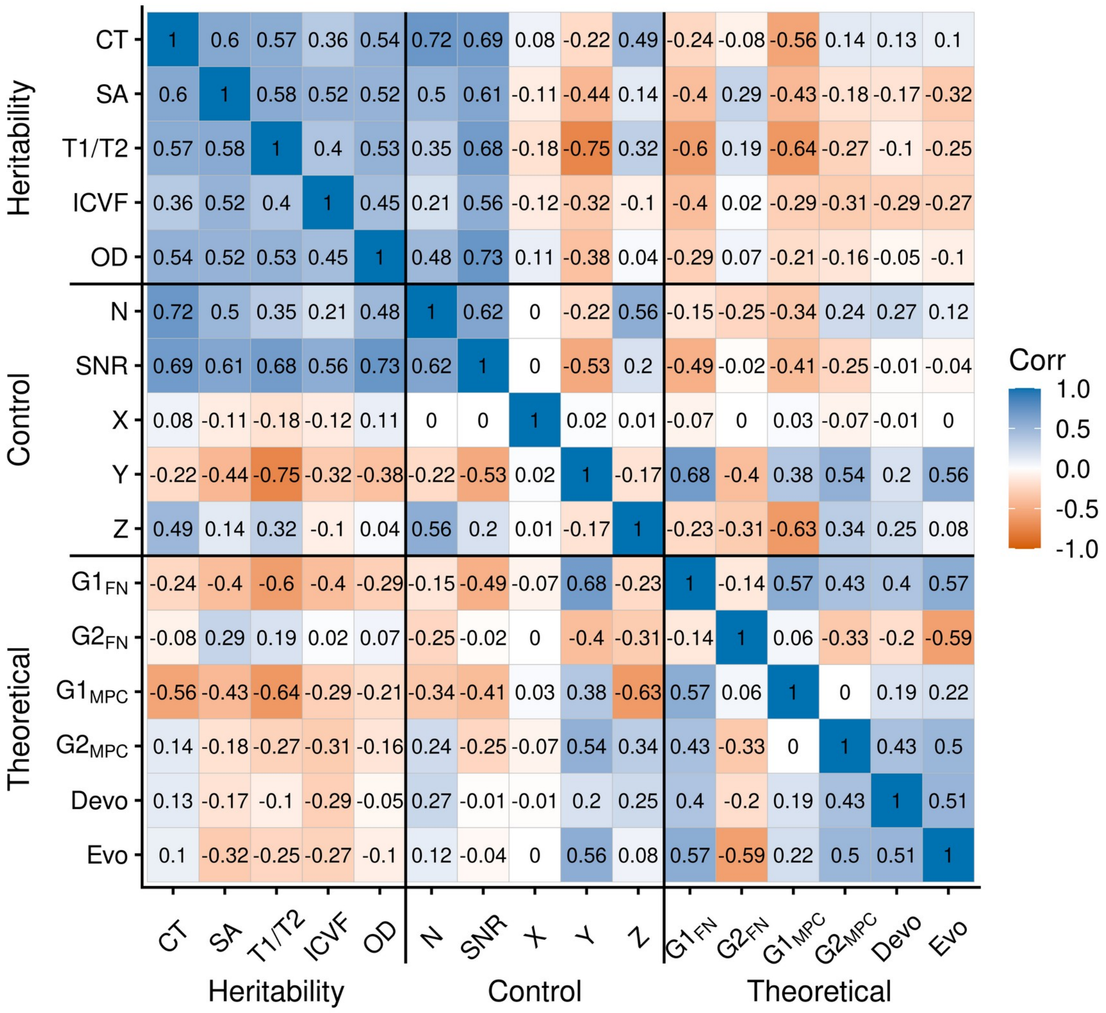


Fig. S2. Spatial correlations between brain maps of heritability across five modalities, brain maps of control variables, and brain maps of explaining variables, using the DKT parcellation. Abbreviations: Corr – correlation; CT – cortical thickness; Devo – developmental; Evo – evolutionary; $G1_{FN}$ – first functional gradient; $G2_{FN}$ – second functional gradient; $G1_{MPC}$ – first structural gradient; $G2_{MPC}$ – second structural gradient; ICVF – intra-cellular volume fraction; N vertices – number of vertices in a parcel; OD – orientation dispersion; SA – surface area; SNR – average signal to noise ratio across five modalities; T1/T2 – T1w over T2w ratio (myelination); X,Y, Z – axis coordinates on the cortex.

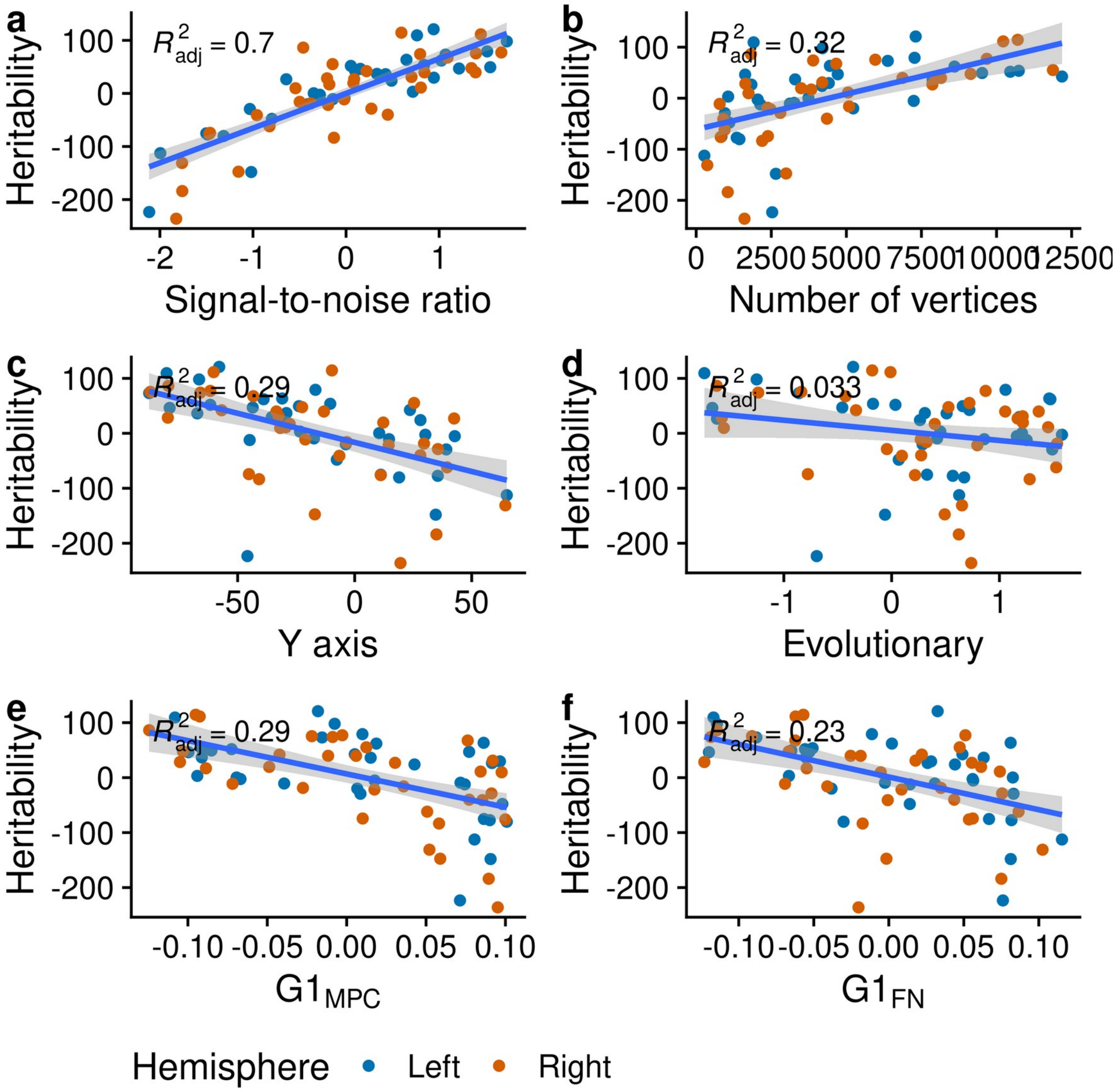


Fig. S3. Scatterplots between mean heritability of DKT parcels and key control and explanatory variables. Abbreviations: $G1_{FN}$ – first functional gradient; $G1_{MPC}$ – first structural gradient.

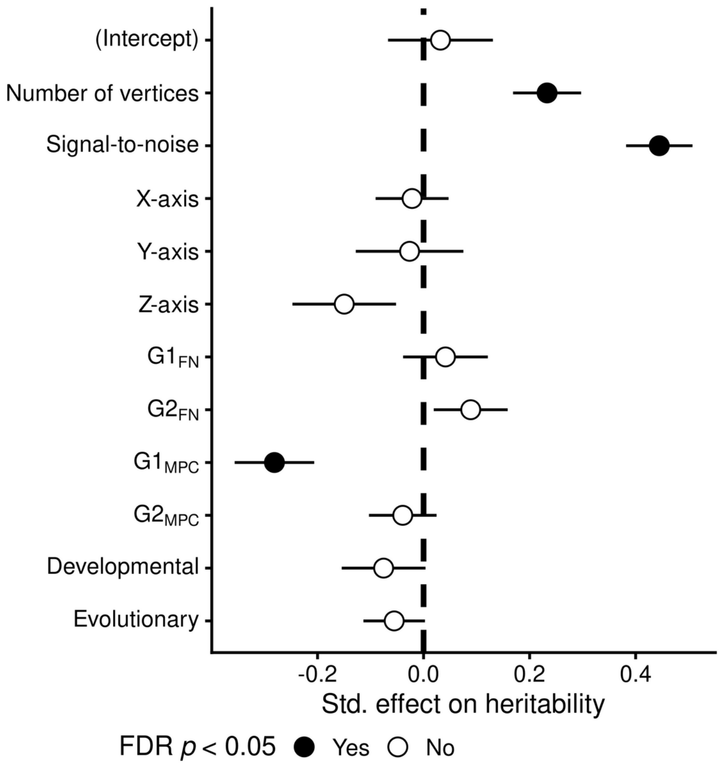


Fig. S4. Independent effects of variables explaining heritability, based on DKT. XYZ are presented, but their effect is zero, as we account for spatial autocorrelation structure. Abbreviations: $G1_{FN}$ – first functional gradient; $G2_{FN}$ – second functional gradient; $G1_{MPC}$ – first structural gradient; $G2_{MPC}$ – second structural gradient.

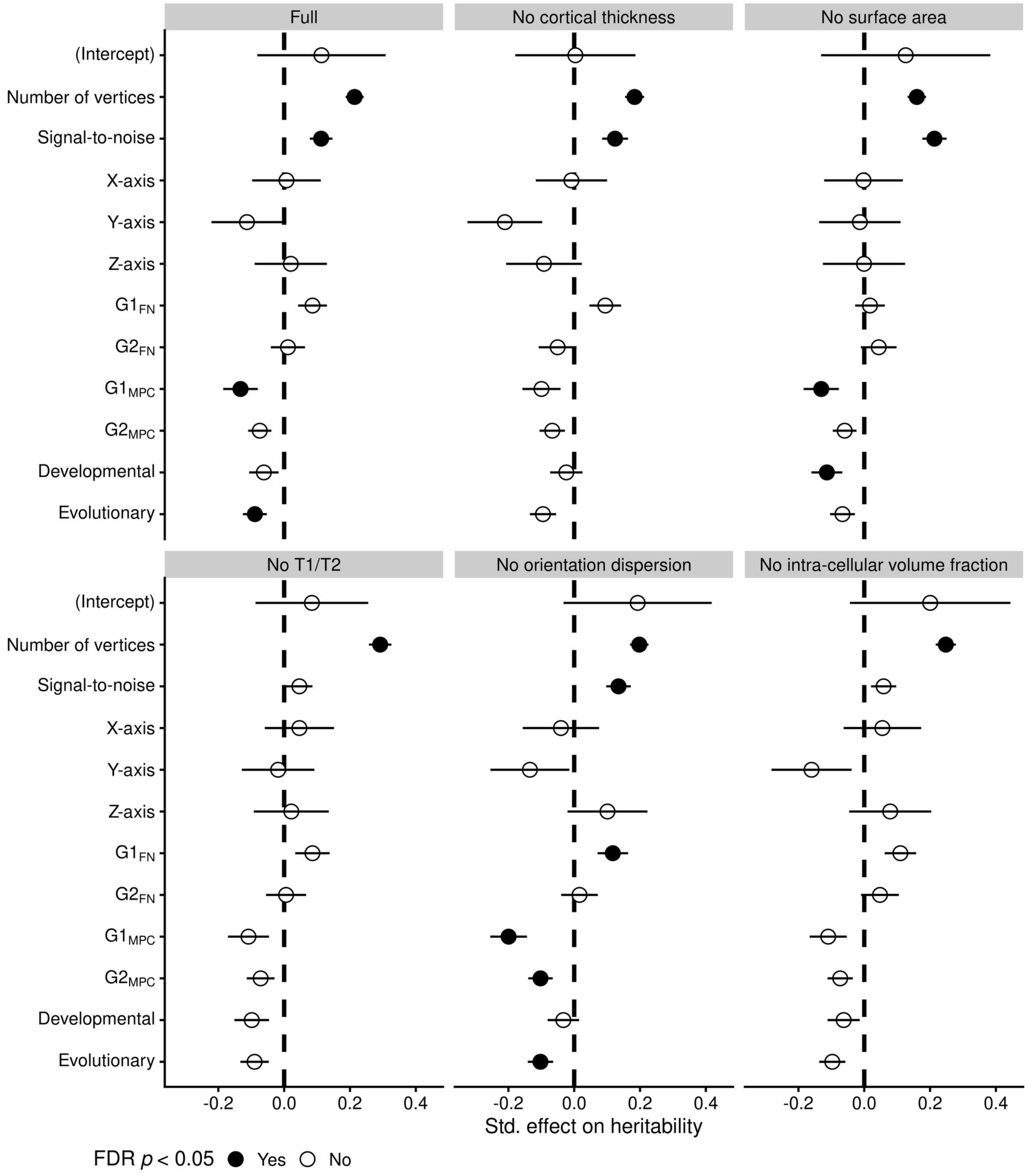


Fig. S5. Leave-one-modality-out replications of the final model, using Schaefer 200 parcellation. Full model corresponds to the same values as in Fig 4. Abbreviations: $G1_{FN}$ – first functional gradient; $G2_{FN}$ – second functional gradient; $G1_{MPC}$ – first structural gradient; $G2_{MPC}$ – second structural gradient.

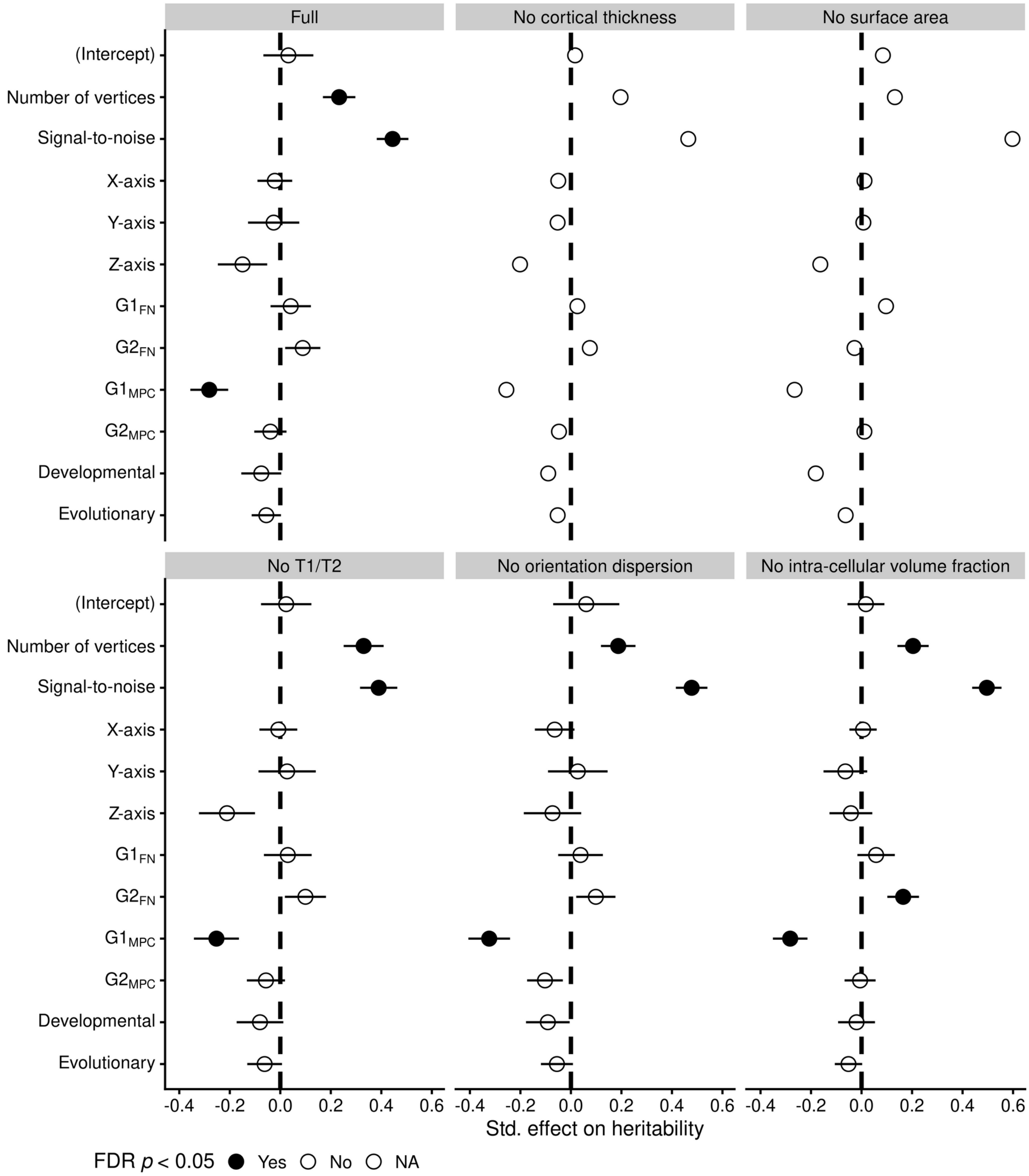


Fig. S6. Leave-one-modality-out replications of the final model, using DKT parcellation. Full model corresponds to the same values as in Fig S4. Abbreviations: $G1_{FN}$ – first functional gradient; $G2_{FN}$ – second functional gradient; $G1_{MPC}$ – first structural gradient; $G2_{MPC}$ – second structural gradient-

Table S1. Summary of models tested.

| Model nr | Model content | Df | AIC | BIC | R2marginal |
|---|---|---|---|---|---|
| M0 | Null model | 3 | 965.8 | 977.3 | |
| M.auto | M0+ exponential autocorrelation | 4 | 831.9 | 847.2 | |
| M.cross.auto | M.auto+crossed random effects | 6 | 813.9 | 836.8 | |
| M.cross.auto.gau | Like M.cross.auto, but with gaussian autocorrelation | 6 | 815.8 | 838.8 | |
| M.ctl.cross.auto | M.cross.auto + control variables | 11 | 734 | 776.2 | 0.69 |
| M.ctl.cross | Like M.ctl.cross.auto, but without autocorrelation | 9 | 766.6 | 801.1 | 0.46 |
| M.ctl.expl.cross.auto | M.ctl.cross.auto + predictors | 17 | 718.8 | 783.9 | 0.70 |

Note. AIC = Akaike Information Criterion. BIC = Bayes Information Criterion. The formula for final glmmTMB model (M.ctl.pred.cross.auto) was Heritability ~control variables + explanatory variables + exp( position + 0 | modality)+ exp( position + 0 | parc), and is summarised in Fig. 4. Marginal R2 characterises the R2 of fixed effects only (Nakagawa et al., 2017).
